# Cyclodextrins increase membrane tension and are universal activators of mechanosensitive channels

**DOI:** 10.1101/2021.03.08.434340

**Authors:** Charles D Cox, Yixiao Zhang, Zijing Zhou, Thomas Walz, Boris Martinac

## Abstract

The bacterial mechanosensitive channel of small conductance, MscS, has been extensively studied to understand how mechanical forces are converted into the conformational changes that underlie mechanosensitive (MS) channel gating. We showed that lipid removal by β-cyclodextrin can mimic membrane tension. Here, we show that all cyclodextrins (CDs) can activate reconstituted *E. coli* MscS, that MscS activation by CDs depends on CD-mediated lipid removal, and that the CD amount required to gate MscS scales with the channel’s sensitivity to membrane tension. CD-mediated lipid removal ultimately causes MscS desensitization, which we show is affected by the lipid environment. CDs can also activate the structurally unrelated MscL. While many MS channels respond to membrane forces, generalized by the ‘force-from-lipids’ principle, their different molecular architectures suggest that they use unique ways to convert mechanical forces into conformational changes. CDs emerge as a universal tool for the structural and functional characterization of unrelated MS channels.

## Introduction

Bacterial mechanosensitive (MS) channels have been extensively used as models of ion channel-mediated mechanotransduction (Cox et al., 2018, Cox et al., 2019). They have continually provided novel insights into the biophysical principles that govern ion-channel mechanosensitivity (Perozo et al., 2002b, Blount and Iscla, 2020, Pliotas et al., 2015, Martinac et al., 1990). While the structurally unrelated MS channels MscL (Blount and Iscla, 2020) and MscS (Booth and Blount, 2012) both respond to changes in membrane tension (Sukharev et al., 1994, Moe and Blount, 2005, Sukharev, 2002), at the molecular level they seem to employ different strategies to convert membrane forces into the conformational changes that underlie channel gating.

*E. coli* MscS is the archetypal member of a large structurally diverse family of ion channels that are expressed in bacteria (Rowe et al., 2013, Martinac et al., 1987), archaea (Kloda and Martinac, 2001), fungi (Nakayama et al., 2012), plants (Haswell and Meyerowitz, 2006) and eukaryotic parasites (Cox et al., 2015). This channel gates as a result of membrane tension (Sukharev, 2002) in accordance with the ‘force-from-lipids’ gating mechanism (Martinac et al., 1990). In response to increases in membrane tension, MscS exhibits complex adaptive gating kinetics (Koprowski et al., 2011, Grajkowski et al., 2005, Akitake et al., 2005). These kinetic responses may represent two separable processes, adaptation and inactivation (Rowe et al., 2014, Kamaraju et al., 2011). In particular, point mutations within transmembrane domain 3 can instigate phenotypes in which adaptation and inactivation are affected differently (Akitake et al., 2007). These complex kinetics are important for the role of this channel as an osmotic safety valve (Boer et al., 2011). However, since it is currently unknown whether these electrophysiologically separable processes correlate to structurally distinct states, we will refer to them collectively as ‘desensitization’. In addition, while some data suggests that MscS desensitization is sensitive to the lipid environment (Xue et al., 2020), this notion still awaits definitive proof.

MscL was the first MS channel to be cloned and functionally characterized in a lipid-only environment (Sukharev et al., 1994). Members of the MscL family, unlike those of the MscS family, are almost exclusively expressed in archaea and bacteria. After X-ray crystallography revealed the structure of MscL (Chang et al., 1998), subsequent studies implicated membrane thinning in response to membrane tension as a major driver of MscL gating (Perozo et al., 2002b).

To fully understand the structural basis of the gating transitions in MscL and MscS, one must first find a way to apply a gating stimulus to the channels in a lipidic environment that is compatible with structural studies. This is, of course, less challenging when considering ligand-gated channels (Hite et al., 2017, Hite and MacKinnon, 2017, Kumar et al., 2020), for which the stimulus is a defined molecule that can readily be applied to visualize the resulting changes in protein conformation. For MS channels, only spectroscopic approaches, such as electron paramagnetic resonance spectroscopy (Vasquez et al., 2008, Perozo et al., 2002a) and Förster resonance energy transfer spectroscopy (Corry et al., 2010, Wang et al., 2014), were available until recently to provide structural insights into their gating in response to changes in forces in their lipid environment. Other approaches had been confined to the use of activators (Brohawn et al., 2014a) or mutations (Wang et al., 2008, Deng et al., 2020). We recently demonstrated that lipid removal by β-cyclodextrin can mimic membrane tension in membrane-scaffold protein-based lipid nanodiscs, providing novel insights into the structural rearrangements that underlie MscS channel gating in response to membrane tension (Zhang et al., 2021).

Cyclodextrins (CDs) are a family of cyclic glucose oligomers with a cone-like 3D architecture characterized by a polar external surface and a hydrophobic cavity (Crini, 2014, Connors, 1997). α, β and γCD contain six, seven or eight glucose units, respectively. As the number of units increases so does the diameter of the hydrophobic cavity (5 to 8 Å) (Connors, 1997). These compounds are of very broad utility as the hydrophobic cavity can chelate a plethora of small lipophilic molecules (Braga, 2019, Carneiro et al., 2019). The hydrophobic cavity of CDs can also form complexes with fatty acids and phospholipids (Szente and Fenyvesi, 2017). CDs have thus been used to remove lipids from native (Vahedi et al., 2020, Startek et al., 2019) and model membranes (Sanchez et al., 2011, Denz et al., 2016), a process that has already been linked to increases in membrane tension (Biswas et al., 2019). CDs also exhibit differential lipid selectivity. For example, αCD has the selectivity profile of phosphatidylserine > phosphatidylethanolamine >> sphingomyelin > phosphatidylcholine (Debouzy et al., 1998). In addition, the methylated version of βCD (mβCD) shows selectivity toward cholesterol at low concentrations and has been widely used in biological research to selectively remove or add cholesterol to cell membranes (Mahammad and Parmryd, 2015, Ridone et al., 2020, Startek et al., 2019, Zidovetzki and Levitan, 2007). In addition to the headgroup, CDs also preferentially chelate unsaturated lipids and those containing shorter acyl chains (Huang and London, 2013, Ikeda et al., 2016).

Here, we show that all members of the CD family (α, β and γ) can activate *E. coli* MscS in liposomal membranes. Even the methylated version of βCD, which is widely used for its cholesterol selectivity, can activate *E. coli* MscS. The CD amount required for the activation of a channel depends on its tension sensitivity. Our studies also clearly establish that MscS desensitization is modified by the lipid environment. Moreover, we show that CD-mediated lipid removal causes a concentration- and time-dependent increase in the tension in membrane patches and that the resulting tension can become sufficiently high to activate the structurally unrelated MS channel MscL that gates at membrane tensions immediately below the lytic limit of membranes. 2D-class averages of nanodisc-embedded MscL obtained by cryo-electron microscopy indicate that βCD treatment results in membrane thinning and channel expansion. The fact that CD activates MscL, which opens immediately below the lytic limit of the membrane, suggests that all other MS channels (which are more sensitive to membrane tension) should also open in response to CDs. These data suggest that CDs will be of broad utility for the structural and functional characterization of structurally diverse MS channels, including Piezo channels (Cox et al., 2016, Syeda et al., 2016), two-pore domain K^+^ channels (Brohawn et al., 2014b, Clausen et al., 2017) and OSCA channels (Murthy et al., 2018, Zhang et al., 2018), all of which are known to respond to membrane forces.

## Results

### All cyclodextrins activate MscS reconstituted in azolectin liposomes

We first tested whether all cyclodextrins (CDs) could activate wild-type MscS. For this purpose, we purified N-terminally 6xHis-tagged MscS (6xHis-MscS) and reconstituted it into azolectin liposomes using the dehydration-rehydration method. The electrophysiological properties of 6xHis-MscS have been reported to differ from those of C-terminally His-tagged or untagged MscS (Reddy et al., 2019). Nevertheless, 6xHis-MscS reconstituted in azolectin liposomes produced stretch-evoked currents with a sensitivity similar to that of MscS after removal of the N-terminal 6xHis tag. However, since 6xHis-MscS displayed pronounced irreversible desensitization behaviour (SI Fig. 1A-C), the N-terminal 6xHis tag was cleaved off for all further experiments.

The application of α, β, and γCD (10 mM) to excised liposome patches activated the incorporated MscS in the absence of any applied hydrostatic pressure (Fig. 1A-D). Activation occurred rapidly with all three CDs, namely within 45 s of perfusion (combined n = 14). We observed little difference in the time course of activation elicited by the different CDs and with all CDs we observed instances of channel desensitization prior to membrane rupture (Fig. 1C).

**Figure 1.**
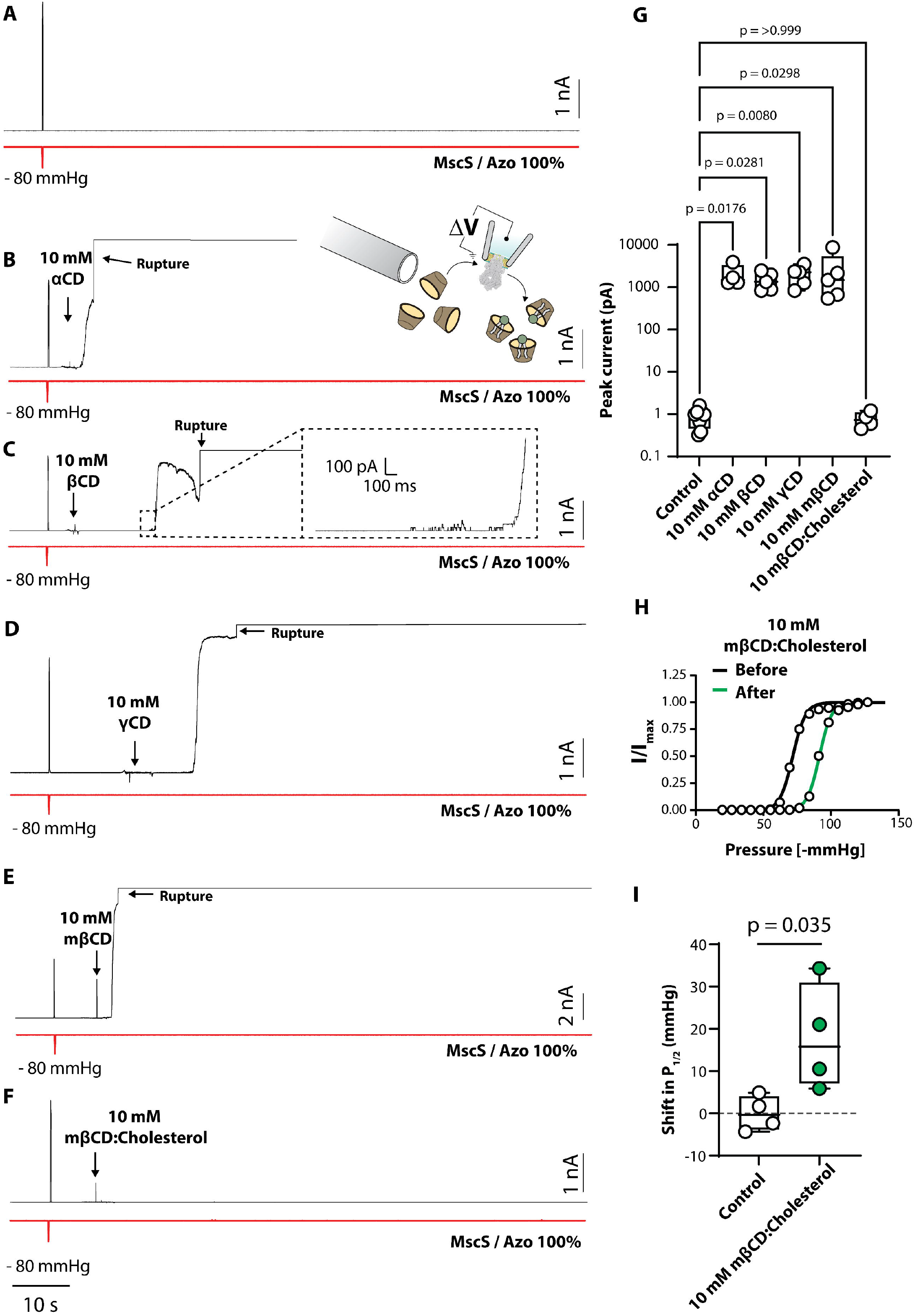
Cyclodextrins (CDs) activate MscS reconstituted into azolectin bilayers. (A) Representative control patch-clamp recording of MscS reconstituted in azolectin (Azo) liposomes (1:200 protein:lipid ratio) at +20 mV pipette potential. The red trace shows the application of negative hydrostatic pressure with a 100-ms ramp up to -80 mmHg. (B-F) Representative patch-clamp recordings of MscS reconstituted in azolectin liposomes in response to perfusion of 10 mM α-cyclodextrin (αCD) (B), 10 mM β-cyclodextrin (βCD) (C), 10 mM γ-cyclodextrin (γCD) (D), 10 mM methylβ-cyclodextrin (mβCD) (E), and 10 mM mβCD loaded with cholesterol (mβCD:cholesterol) (F). These recordings show that all CDs and their derivatives can activate MscS in azolectin liposomes. Note that for all CDs, some traces showed evidence for MscS desensitization, as clearly seen in panel C. (G) Quantification of the peak current elicited from excised liposome patches in response to CD perfusion prior to patch rupture. p-values shown were generated by comparison to the control group (not perfused) according to the Kruskal Wallis test with Dunn’s post hoc. (H) Representative Boltzmann fit of MscS pressure response before (black) and after (green) the perfusion of 10 mM mβCD:cholesterol. The right shift in the pressure response curve denotes the channel is less sensitive to applied force. (I) Quantification of the degree of rightward shift in the pressure response curve of MscS patches not perfused with mβCD:cholesterol (control) compared with those perfused with 10 mM mβCD:cholesterol (T-test used for comparison). Data are represented as minimum to maximum box and whisker plots showing all data points.

Activation also occurred with the methylated derivative of βCD (10 mM mβCD), which is often used to extract cholesterol from membranes (Fig. 1E). In contrast, application of the same concentration of mβCD (10 mM) that was saturated with cholesterol did not induce MscS gating (Fig. 1E-G). In fact, perfusion of cholesterol-saturated mβCD reduced the sensitivity of MscS as evidenced by a rightward shift in the pressure-response curve that was recorded 3 min after the cholesterol-saturated mβCD was added (Fig. 1G-H). This observation suggests that cholesterol was transferred from mβCD into the membrane patch, as cholesterol has previously been shown to reduce the mechanosensitivity of *E. coli* MscS (Nomura et al., 2012). Together, these results strongly support the notion that CDs activate MscS by creating membrane tension as a result of removing lipids from the membrane, whereas inserting additional lipids, in particular cholesterol that further reduces membrane fluidity and increases membrane stiffness by intercalating in between the lipid acyl chains, creates membrane pressure that inhibits MscS.

Addition of CDs also resulted in prolonged sojourns of MscS into sub-conducting states, mirroring results previously obtained by applying sustained mechanical tension on MscS in azolectin patches, which also resulted in MscS assuming numerous sub-states (SI Fig. 2A-C) (Cox et al., 2013). This result therefore implies that CD-mediated lipid removal results in sustained membrane tension. Recently, we proposed that the structure of MscS in the open state is dynamic (Zhang et al., 2021) and the multiple sub-states observed here support this notion.

### Cyclodextrin removes lipids from excised liposome patches

To further corroborate that MscS activation by CDs is due to lipid removal, we perfused membrane patches with CD and concomitantly recorded MscS activation and imaged the patch membrane. The fluorescent lipid rhodamine-PE18:1 was added to both visualize the patch and monitor the lipid content. In empty liposomes with no channel protein incorporated, perfusion with 5 or 10 mM βCD caused a graded reduction in fluorescence intensity within the patch (Fig. 2A). No quantal events signifying channel openings were observed up until the patch ruptured. In patches containing MscS, we observed that MscS activity occurred concomitantly with a reduction in the dome height of the liposome patch (Fig. 2B-D). This flattening of the dome was evident when the dome height was measured over time (Fig. 2C). The resting inward curvature seen at the beginning of the recordings is only possible in the presence of excess lipids. As the lipids were removed, the patch membrane flattened, and this coincided with channel activation (Fig. 2B).

**Figure 2.**
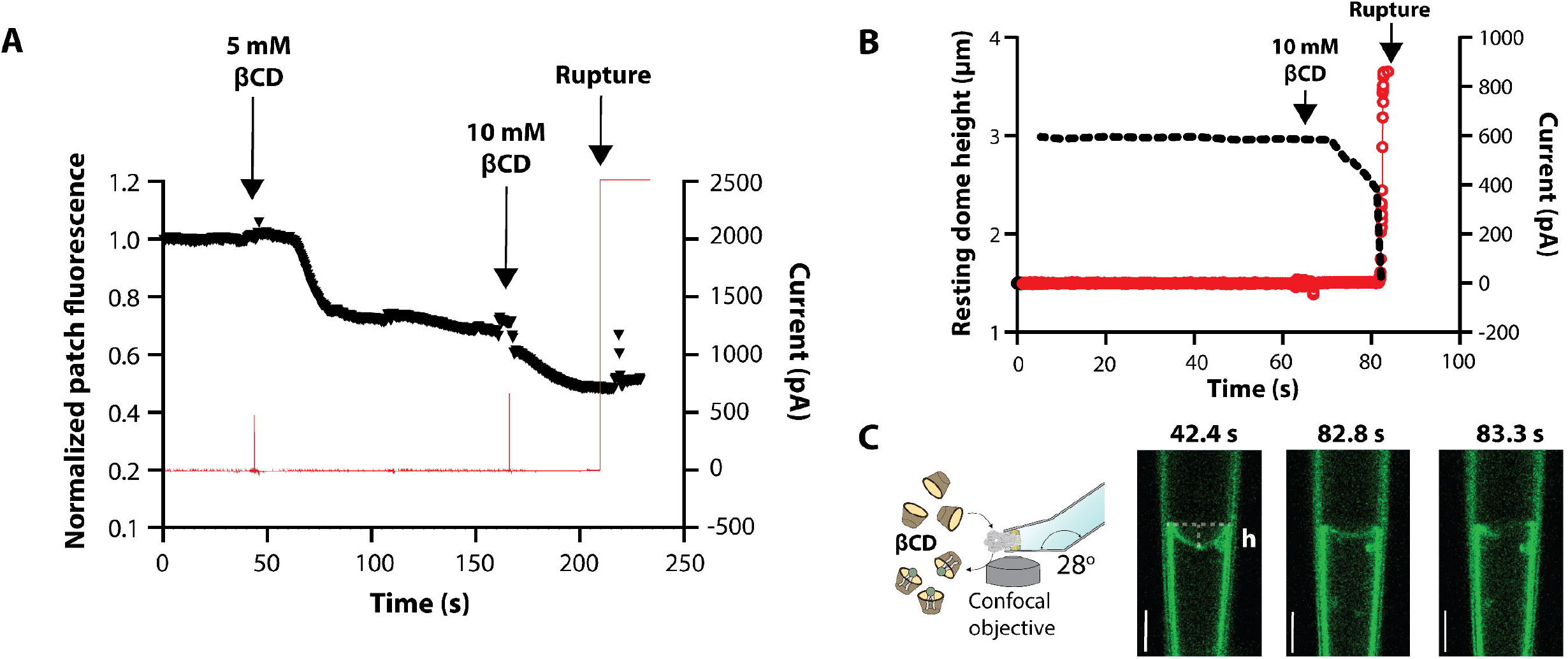
β-cyclodextrin removes lipids from excised membrane patches. (A) Normalized patch fluorescence of an azolectin liposome patch containing 0.1% rhodamine-PE (black) without any reconstituted channels. Red trace shows concomitant electrophysiology trace showing no channel activity and ultimate patch rupture. (B) Height of the patch dome measured concomitantly with electrophysiological recording of the activity of MscS reconstituted in azolectin liposomes supplemented with 0.1% rhodamine-PE. Black trace shows how the dome height changes with time and red trace represents channel current. Time point when 10 mM βCD was added is indicated. (C) Cartoon illustrating the experimental set up (left) with representative confocal images of the patch dome during patch-clamp recordings at indicated time points (right). h denotes the height of the dome. White vertical scale bars, 5 µm.

### The amount of cyclodextrin required to activate MscS scales with its sensitivity to membrane tension

The tension sensitivity of MscS is affected not only by mutations but also by the lipid composition of the surrounding membrane (Xue et al., 2020, Nomura et al., 2012). We recently reported that doping azolectin liposomes with 30% (w/w) PC10, a lipid with two short 10-carbon acyl chains, causes spontaneous short-lived sub-state openings of MscS (Zhang et al., 2021). While electrophysiological (Nomura et al., 2012, Xue et al., 2020) and structural data (Zhang et al., 2021) suggest hydrophobic mismatch is not the main driver of MscS channel gating, we found that as little as 15% PC10 does increase the tension sensitivity of MscS (Fig. 3A-C). In addition, we observed that PC10 substantially slowed channel closure, with MscS remaining open for seconds after the applied hydrostatic pressure had returned to zero (Fig. 3A, D).

**Figure 3.**
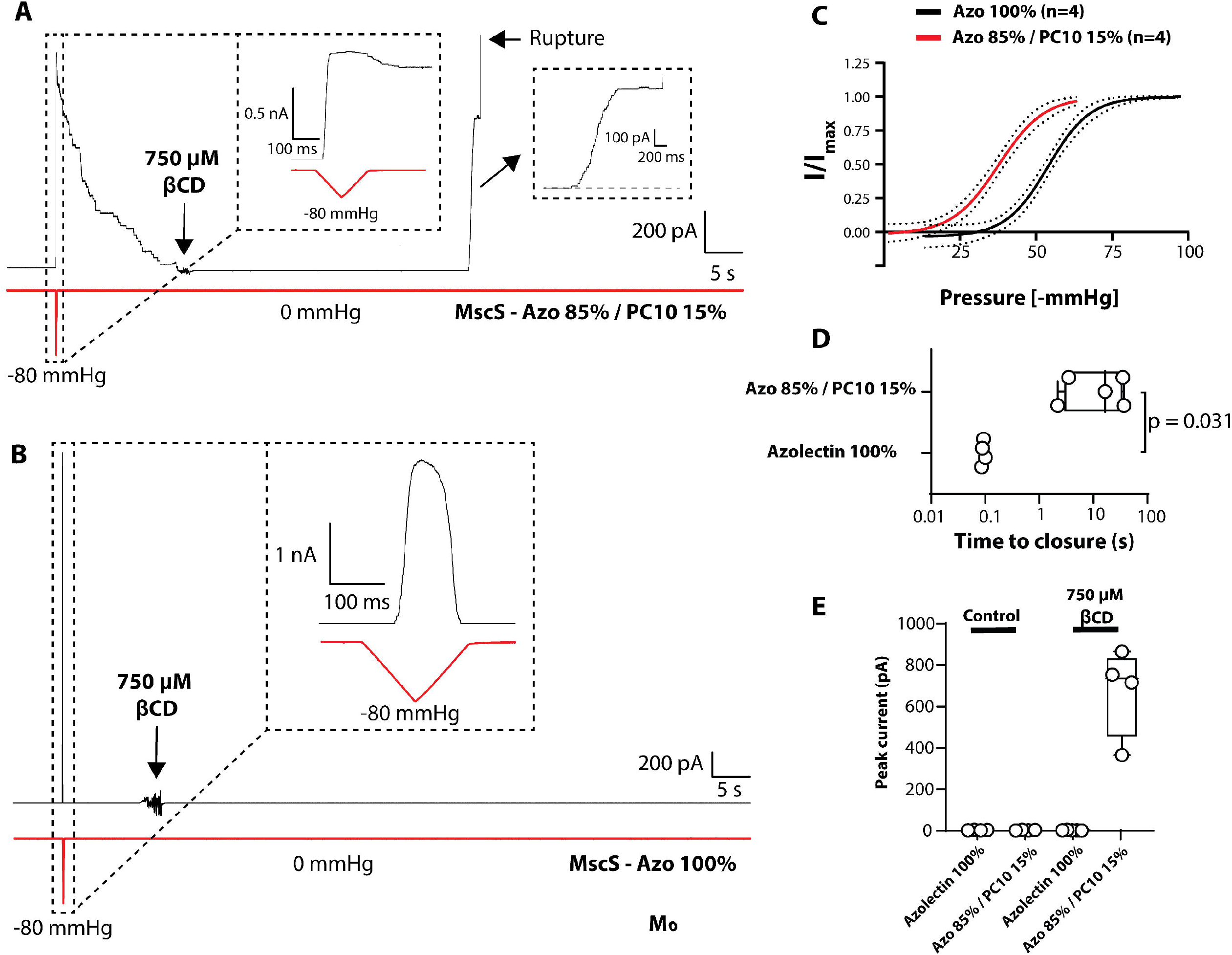
The cyclodextrin concentration needed to induce gating scales with the tension sensitivity of MscS. (A) Representative patch-clamp recording of MscS reconstituted in azolectin (Azo) liposomes containing 15% PC10 (1:200 protein:lipid ratio) at +20 mV pipette potential with the addition of 750 µM βCD. (B) Representative patch-clamp recording of MscS reconstituted in pure azolectin liposomes (1:200 protein:lipid ratio) at +20 mV pipette potential with the addition of 750 µM βCD. (C) Comparison of the pressure-response curves of MscS reconstituted in azolectin alone (black) and in azolectin containing 15% PC10 (red). Data are fitted with Boltzmann distribution and dashed black lines show the 95% confidence interval. (D) Quantification of the time from peak pressure applied until the last MscS closes in pure azolectin liposomes and azolectin liposomes containing 15% PC10. (E) Quantification of the peak current elicited from excised liposome patches in response to perfusion with 750 µM βCD prior to patch rupture. These results illustrate that lower amounts of βCD are required to activate MscS in a lipid environment in which the required tension for channel gating is lower.

We hypothesized that in lipid environments in which MscS is more sensitive to membrane tension fewer lipids would need to be removed by CDs to generate sufficient force to open the channel. To test this hypothesis, we compared the effect of much lower amounts of βCD (<10 mM) on MscS activity in liposomes containing 15% PC10. In such a membrane environment, MscS was indeed activated by βCD concentrations as low as 750 µM (Fig. 3A, D). In comparison, over the same electrophysiological recording period, 750 µM βCD failed to elicit any gating events for MscS that was incorporated in azolectin liposomes that contained no PC10.

### Cyclodextrin-induced activation reveals that MscS desensitization is affected by the lipid environment

As shown in Figure 1C, βCD induces MscS activation followed by desensitization. *E. coli* membranes are predominantly composed of phosphatidylethanolamine (PE) lipids, so to more closely mimic the native lipid environment (Bogdanov et al., 2020) of the channel and to examine the effect of membrane composition on MscS desensitization, we reconstituted MscS into pure azolectin liposomes and liposomes made of 70% azolectin with 30% PE18:1. Incorporation of MscS in liposomes containing 70% azolectin and 30% PE18:1 right-shifted the pressure response of MscS (P_1/2_ = 71.1; 95% confidence interval: 70-74.5) compared to channels reconstituted in pure azolectin liposomes (P_1/2_ = 53.9; 95% confidence interval: 51.5-56.1). Figure 3 shows that lower amounts of βCD were required to generate MscS activity when the channel was reconstituted in PC10-containing liposomes, in which case lower tensions sufficed to open MscS. Consistent with this result, more βCD was required (i.e. 15 mM) to elicit gating events in liposomes doped with 30% PE18:1 (Fig. 4A), in which case higher tensions are needed to open MscS. Lower βCD amounts failed to elicit MscS activity in liposomes with 30% PE18:1 (0/6 patches with 5 mM βCD) over the same recording period (90 s). Moreover, after activation of MscS in liposomes doped with 30% PE18:1, we observed rapid full desensitization of the channel (Fig. 4A). Conversely, Gly113Ala mutant MscS, which displays abrogated desensitization (Akitake et al., 2007, Edwards et al., 2008), was activated by lower concentrations of βCD (e.g. 5 mM), even in liposomes containing 30% PE18:1. This mutant stayed open for long periods (>30 s) prior to patch rupture, with little evidence of desensitization (Fig. 4B). Importantly, for the Gly113Ala mutant channel in liposomes containing 30% PE18:1, the time from activation of the first channel to maximal current is substantially longer (23 ± 12 s; n = 5) when 5 mM βCD was used than when 10 mM CD was used as shown in Figure 1 (3.4 ± 1.9 s; n = 14). These results suggest that MscS desensitization kinetics depend on the lipid environment.

**Figure 4.**
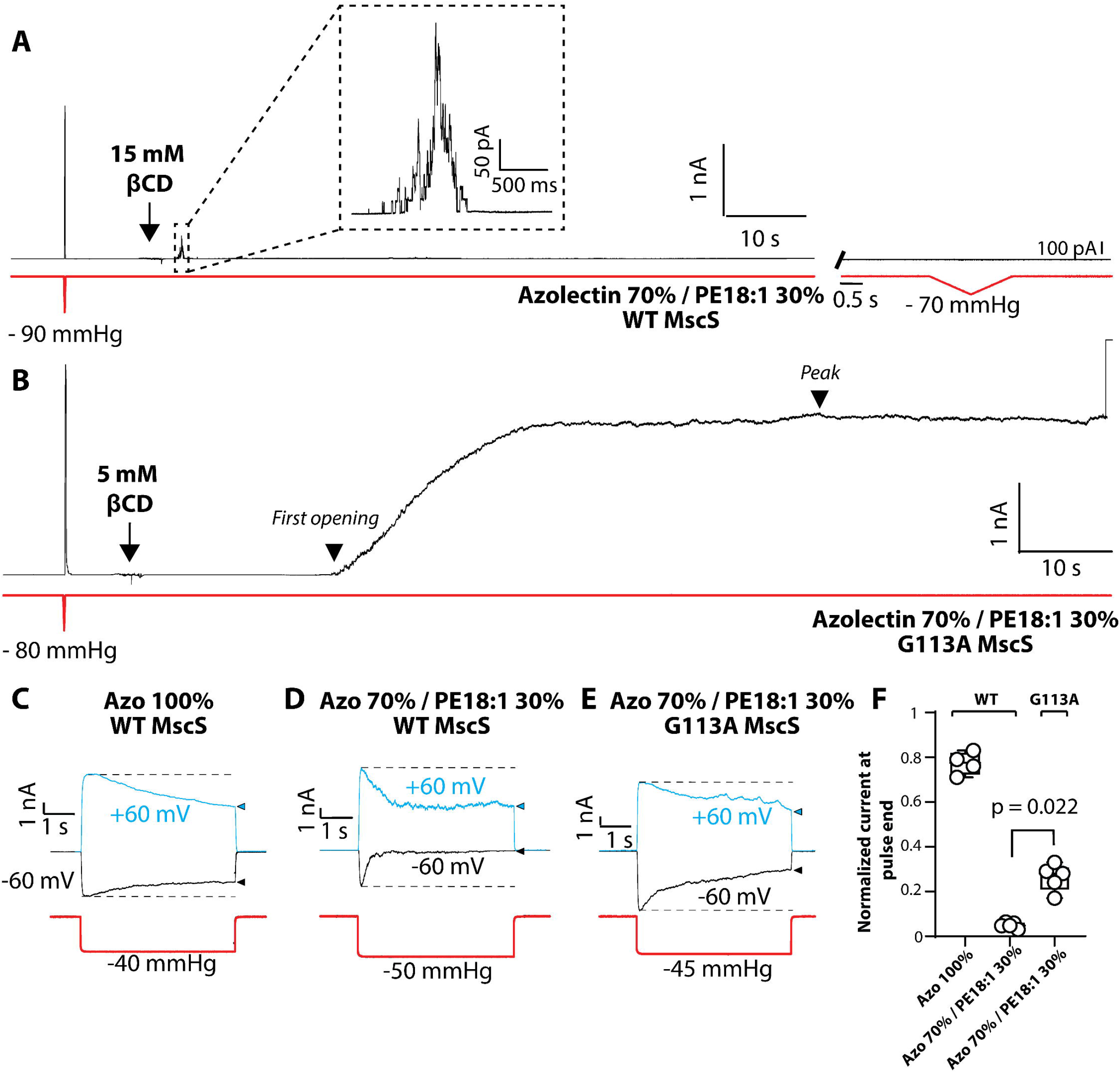
Cyclodextrins induce activation and subsequent desensitization of MscS, which is influenced by the lipid environment. (A) Representative electrophysiological recordings of wild-type (WT) MscS reconstituted in 70% azolectin (Azo) and 30% PE18:1 at +20 mV pipette potential in response to perfusion with 15 mM βCD. After activation, the channels rapidly desensitize (left) and then remain insensitive to applied negative pressure (right). (B) Representative electrophysiological recordings of Gly113Ala mutant (G113A) MscS, a channel variant that does not desensitize, reconstituted in 70% azolectin and 30% PE18:1 at +20 mV pipette potential in response to perfusion with 5 mM βCD. Once the mutant channels are activated by lipid removal, they stay open for a long period with little to no signs of desensitization. (C) Representative electrophysiological recordings of WT MscS reconstituted in 100% azolectin at pipette potentials of +60 mV (blue) and -60 mV (black). (D) Representative electrophysiological recordings of WT MscS reconstituted in 70% azolectin and 30% PE18:1 at pipette potentials of +60 mV (blue) and -60 mV (black) showing rapid desensitization at negative potentials. (E) Representative electrophysiological recordings of Gly113Ala mutant MscS reconstituted in 70% azolectin and 30% PE18:1 at pipette potentials of +60 mV (blue) and -60 mV (black). Note the much slower desensitization of this mutant channel compared to WT MscS, particularly at -60 mV. (F) Quantification of MscS desensitization as a function of lipid composition at -60 mV pipette potential by comparing the peak current to the current remaining at the end of a 5-s sub-saturating pressure pulse at 40% of the saturating pressure.

To further illustrate this fact, we used 5-s-long sub-saturating pressure pulses at 40% of the saturating pressure to normalize desensitization between patches (Fig. 4C-F) (Cetiner et al., 2018, Grajkowski et al., 2005). Sub-saturating pressure pulses more readily reveal MscS desensitization kinetics. In addition, desensitization is voltage-dependent and is more readily observed at negative pipette potentials than at positive pipette potentials, shown in Figure 4C-E by comparing channel activity at +60 mV and -60 mV (Fig. 4C-E). While desensitization is rapid at negative pipette potentials (−60 mV) for wild-type MscS in the presence of 30% PE18:1, it is much slower in pure azolectin liposomes at the same voltage (−60 mV) as quantified by the current remaining at the end of the pressure pulse (Fig. 4A-D). To prove that the changes in MscS activity in an azolectin membrane compared to that in an azolectin membrane doped with 30% PE18:1 are linked to desensitization, we again reconstituted the Gly113Ala mutant MscS that displays a vastly reduced adaptive gating behavior (Akitake et al., 2007, Edwards et al., 2008). At the same voltage (−60 mV), the Gly113Ala mutant MscS showed substantially less desensitization in an azolectin membrane that contains 30% PE18:1 (Fig. 4D-E).

### Cyclodextrins also activate reconstituted MscL

To test whether CD-mediated lipid removal also activates structurally distinct MS channels, we co-reconstituted MscS with MscL in azolectin liposomes and perfused excised patches with βCD. Under the application of a ramp to a maximum pressure of -40 mmHg, only MscS activity was recorded. To measure the membrane tension during our patch-clamp recordings resulting from the application of negative pressure and βCD perfusion, we concomitantly imaged the rhodamine-PE18:1-containing membrane patch by confocal microscopy. Visualization of the patch demonstrated that the maximum pressure applied (−40 mmHg) generated a membrane tension of 7.1 mN/m, which was calculated using LaPlace’s law by measuring the patch dome radius (Nomura et al., 2012). This tension is saturating for MscS in azolectin (Nomura et al., 2015, Nomura et al., 2012, Sukharev, 2002) but is not high enough to open MscL (Nomura et al., 2012) (Fig. 5A). The patch was then perfused with 25 mM βCD. Perfusion with βCD first opened MscS and as the MscS current saturated the first MscL openings became evident (3x the current amplitude of MscS). Subsequently, many more MscL opened prior to patch rupture (Fig. 5B). Monitoring the membrane patch using confocal microscopy showed that perfusion with βCD caused a progressive loss of the resting inward curvature (Fig. 5B; image *i*) until the patch became completely flat (Fig. 5B; image *iv*) and finally ruptured (Fig. 5B; image *v*).

**Figure 5.**
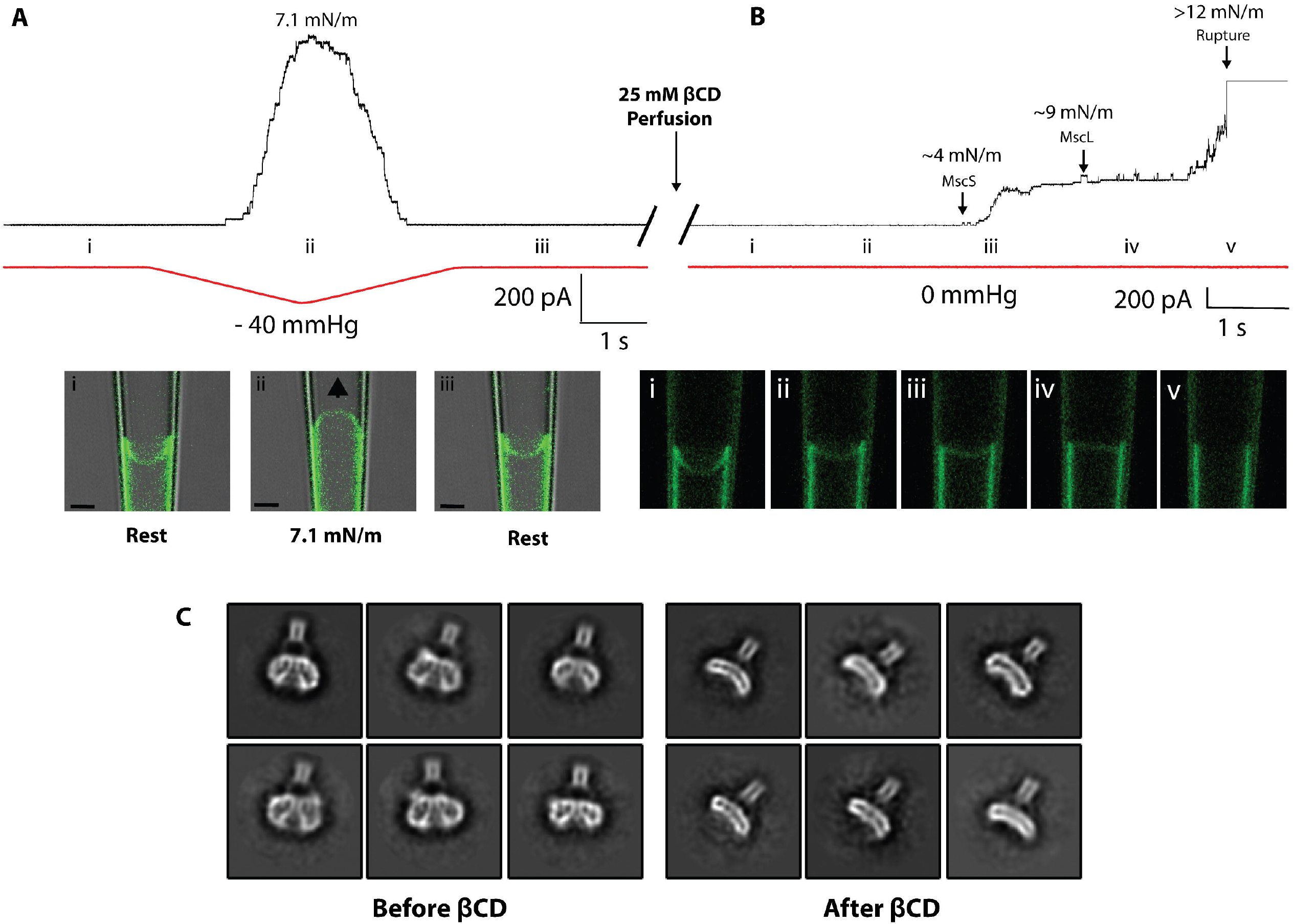
Cyclodextrin treatment activates MscL in liposomes and causes a conformational change in nanodisc-embedded MscL. (A) Top: Exemplar trace of MscS and MscL co-reconstituted in azolectin liposomes (protein:lipid ratios of 1:200 for MscS and 1:1000 for MscL) recorded with a negative pressure ramp to a maximum of -40 mmHg in 1 s. Only MscS activity is seen and no MscL is activated (red trace: pressure; black trace: current). Below: Concomitant confocal imaging of the patch containing 0.1% of the fluorescent lipid rhodamine-PE illustrates the deformation of the membrane patch caused by the negative pressure that allowed calculation of the maximal tension generated (7.1 mN/m). (B) Top: Electrophysiological recording of the same patch after addition of 25 mM βCD, showing first MscS activity (activation threshold ∼5 mN/m), then MscL activity (activation threshold ∼9 mN/m), and finally rupture of the membrane patch (the rupture tension of azolectin patches is >12 mN/m). Below: Concomitant confocal imaging of the membrane patch shows a progressive loss of rhodamine-PE fluorescence and a flattening of the dome prior to rupture. (C) Selected cryo-EM 2D-class averages of nanodisc-embedded MscL before and after βCD treatment (also see SI Figure 3). Note the expansion and thinning of the transmembrane domain. Side length of individual averages, 20 nm.

### βCD-induced conformational change in MscL

Given the fact that βCD could activate MscS in azolectin liposomes and allowed cryo-EM visualization of nanodisc-embedded MscS in the desensitized state, we asked whether βCD-mediated lipid removal would also affect the conformation of MscL. Wild-type MscL was purified and reconstituted into nanodiscs and subsequently treated with 100 mM βCD for 16 h. Cryo-EM imaging and resulting 2D-class averages of the nanodisc-embedded MscL showed that βCD treatment caused an expansion and thinning of its transmembrane domain (Fig. 5C & SI Fig. 3). This observation is consistent with the previously proposed gating mechanism (Perozo et al., 2002a) and structural dynamics (Bavi et al., 2017) of MscL and suggests that βCD-mediated removal of lipids from the nanodiscs resulted in activation of MscL.

### The activating cyclodextrin concentration scales with the tension sensitivity of MscL

To further affirm that the amount of CD necessary to activate an MS channel scales with the tension sensitivity of the channel, we made use of Gly22Ser mutant MscL that has an activation threshold close to that of MscS (Yoshimura et al., 1999). While 5 mM βCD activated multiple Gly22Ser MscL channels over a 90-s recording period, it failed to activate any wild-type MscL channels (Fig. 6A-C). We also confirmed that Gly22Ser mutant MscL reconstituted in azolectin liposomes displays a leftward shift in the pressure-response curve, showing that less membrane tension is required for channel gating (Fig. 6D).

**Figure 6.**
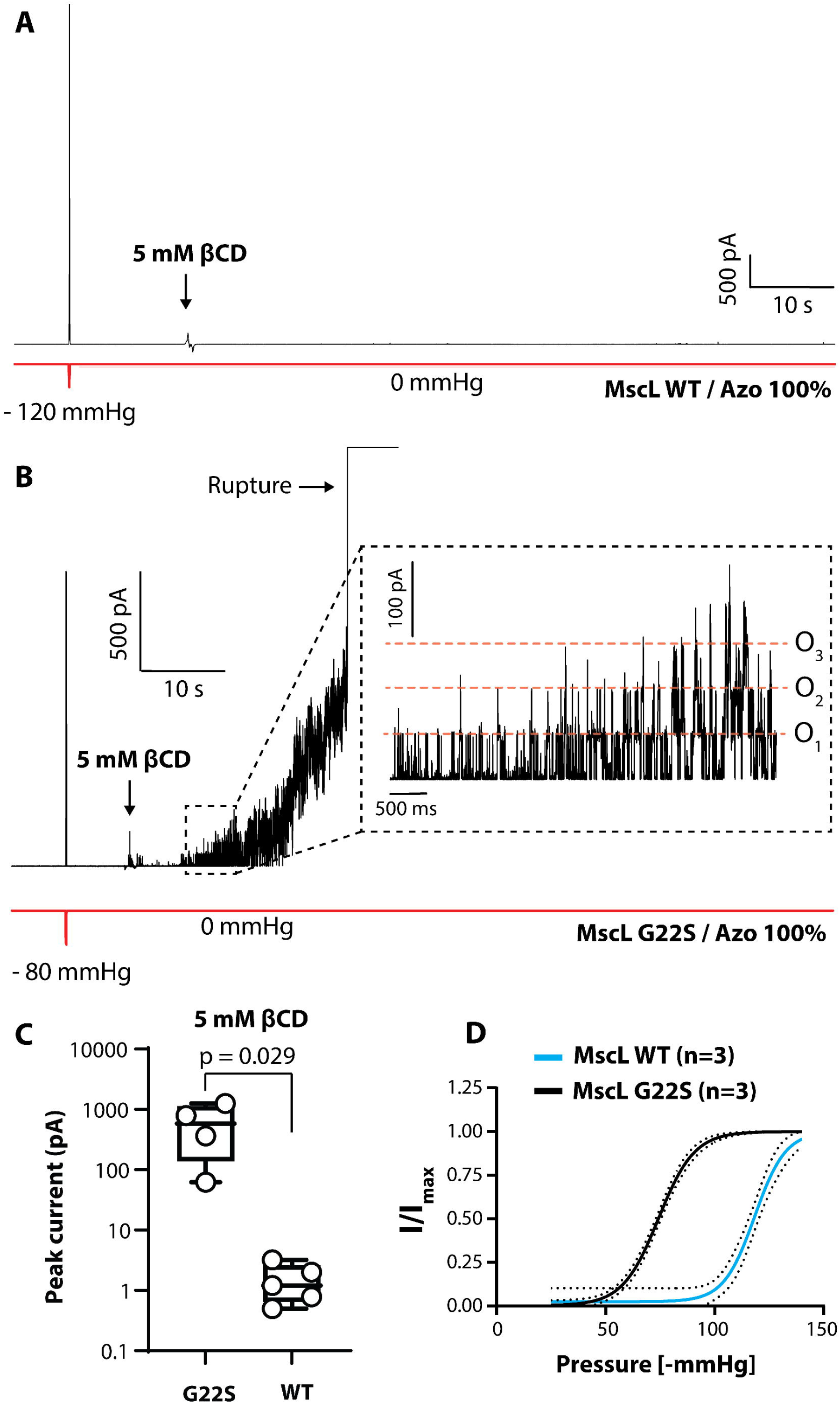
Cyclodextrin concentration needed to activate MscL scales with the tension sensitivity of the channel. (A) Representative patch-clamp recording of wild-type MscL (MscL WT) reconstituted in azolectin (Azo) liposomes (1:1000 protein:lipid ratio) at +20 mV pipette potential with the addition of 5 mM βCD. (B) Representative patch-clamp recording of Gly22Ser MscL (MscL G22S), which gates at a lower membrane tension, reconstituted in azolectin liposomes (1:1000 protein:lipid ratio) at +20 mV pipette potential with the addition of 5 mM βCD (O_1-3_ = channel opening 1-3). (C) Quantification of the MscL peak current elicited from excised liposome patches in response to perfusion with 5 mM βCD for 90 s for WT and G22S MscL. (D) Comparison of the pressure-response curve of WT MscL (blue) and G22S MscL (black) reconstituted in azolectin liposomes. Data are fitted with Boltzmann distribution and dashed black lines show the 95% confidence interval.

## Discussion

MS channels play a key role in many mechanotransduction processes from osmoregulation in bacteria (Naismith and Booth, 2012) to touch and proprioception in mammals (Martinac and Cox, 2017). Structurally diverse classes of MS channels (Jin et al., 2020) have been shown to respond to forces through the membrane according to the force-from-lipids mechanism (Martinac et al., 1990). However, given the diversity in structure and sensitivity between members of MS channel families, it is unlikely that these channels use identical molecular mechanisms to sense membrane perturbations. Here, we show that lipid removal by CDs and their derivatives (mβCD) can induce membrane tension *in vitro* that is sufficient not only to activate MscS but also the structurally unrelated, much less tension-sensitive MscL. This result suggests that CDs will be a useful and generally applicable tool for the functional and structural interrogation of MS channel gating mechanisms.

The fact that the methylated version of βCD can also induce sufficient membrane tension in pure phospholipid membranes is of general relevance. The increase in tension and subsequent activation of MscS by mβCD, which is widely used to extract cholesterol from cell membranes, suggests that experiments using high amounts of mβCD may have unintended effects due to the activation of MS channels (Zidovetzki and Levitan, 2007) and demonstrates a need for appropriate controls when mβCD is used to remove cholesterol from native cell membranes (Zidovetzki and Levitan, 2007).

The mechanism of CD-induced lipid removal likely involves a step of adsorption (Lopez et al., 2011), and adsorption to the bilayer could make CDs activate MS channels in a similar fashion as amphipaths (Cox and Gottlieb, 2019, Martinac et al., 1990). Amphipaths are thought to intercalate or adsorb into the bilayer, thus modifying the local membrane forces and inducing MS channel activation, with examples including lyso-lipids and small molecules like chlorpromazine (Cox and Gottlieb, 2019, Lundbaek et al., 2010, Martinac et al., 1990). However, we have two pieces of evidence that support the conclusion that CDs increase membrane tension *in vitro* by lipid removal and do not act simply as amphipaths. First, cholesterol-loaded mβCD does not instigate MscS gating *in vitro*, even though lipid delivery almost certainly also involves a step in which mβCD adsorbs to the membrane. Second, fluorescence imaging of membrane patches shows that lipid removal, as evidenced by the reduction in its fluorescence and the loss of the patches’ resting inward curvature, happens on the same time scale as MS channel activation.

We also show using two different approaches that the amount of CD necessary to activate an MS channel in a liposomal membrane scales with the sensitivity of the channel to membrane tension. First, we observed that MscS exhibited a left-shifted pressure-response curve when reconstituted in azolectin membranes doped with 15% PC10. The increased tension sensitivity of MscS in this membrane environment resulted in lower concentrations of βCD (<1 mM) being necessary to induce gating. Second, we demonstrated that Gly22Ser mutant MscL that is more sensitive to membrane tension (Yoshimura et al., 1999) could be activated by lower amounts of βCD than wild-type MscL over similar recording periods. These results provide strong evidence that the amount of CD needed to activate an MS channel, and thus the amount of lipid that needs to be removed from the membrane, is proportional to how sensitive the channel is to membrane tension.

The βCD-induced activation of MscS in nanodiscs and membrane patches is followed by desensitization of the channel (Zhang et al., 2021). Structurally the ‘desensitized state’ that βCD induces resembles what has electrophysiologically been characterized as the inactivated state (Kamaraju et al., 2011, Rowe et al., 2014). For example, Asp62 and Arg128 form a salt bridge in the inactivated state (Rowe et al., 2014, Nomura et al., 2008) and this salt bridge is seen in the cryo-EM structure of MscS in the desensitized state. Previous studies of MscS desensitization have shown that MscS is activated most robustly and fully by rapid force application in the form of a steep pressure ramp (Cetiner et al., 2018, Akitake et al., 2005). Slower force application reduces the number of activated channels and promotes desensitization (Akitake et al., 2005, Akitake et al., 2007). Interestingly, larger amounts of βCD are needed to activate MscS in azolectin liposomes doped with 30% PE18:1 compared with MscS in pure azolectin liposomes. This finding is consistent with the previously reported result that higher tension is required to open MscS in this lipid mixture (Xue et al., 2020). Furthermore, channel activation in this lipid mixture is followed by rapid desensitization. Using a normalized pressure protocol at sub-saturating pressures, we showed that MscS desensitization is more prominent in membranes doped with 30% PE18:1 (Xue et al., 2020). A possible explanation for these observations is that lower βCD concentrations result in a slower removal of lipids and thus a slower build-up of membrane tension. This slower increase in membrane tension is more akin to the forces generated by slower pressure ramps. Slower ramps activate some MscS, which then close and desensitize, resulting in much lower peak currents. In fact, in some scenarios, MscS has been shown to ‘silently’ transition from the closed to a desensitized state (without entering the open state) (Belyy et al., 2010), which could also explain why lower βCD concentrations cannot activate MscS in 30% PE18:1 liposomes. This conclusion is supported by the fact that Gly113Ala mutant MscS that displays markedly reduced desensitization can be activated by lower amounts of βCD and even in the presence of 30% PE18:1, where there is a gradual activation of all channels in the patch and no sign of any desensitization. PE lipids are the most abundant lipid type in *E. coli* and increasing PE content within liposomes produces MscS activity that is more similar to that seen with native spheroplast membranes, which is characterized by a higher-pressure threshold and more prominent desensitization (Akitake et al., 2005, Shaikh et al., 2014).

As *in-vitro* studies on MS channels begin to involve more complex lipid mixtures, which mimic the lipid compositions of their native membrane environments, the differential selectivity of CD subtypes may provide some control over which lipids are removed from the membrane to induce membrane tension (Huang and London, 2013). It may be particularly important not to remove lipids that are critical for the function or mechanosensitivity of MS channels, which applies not only to structural studies on MS channels reconstituted in nanodiscs but also to functional studies, such as patch-clamp electrophysiology, planar bilayers (Clausen et al., 2017) and liposome flux assays (Cabanos et al., 2017, Su et al., 2016, Mukherjee et al., 2014). Here, we should note that it is not only MS channels that are sensitive to mechanical forces, and CDs may thus also prove useful for the *in-vitro* study of other membrane proteins that are sensitive to mechanical forces (Kim et al., 2020, Xu et al., 2018, Erdogmus et al., 2019).

In conclusion, we present an extensive characterization of CD-induced activation of MscS *in vitro* in patch-clamp experiments. The three major classes of CDs, including a methylated derivative, can all rapidly activate MscS, consistent with a rise in membrane tension resulting from lipid extraction. We also show that CDs not only activate MscS but that higher amounts perfused over the same timescale can also rapidly activate the structurally unrelated MscL. This result suggests that for both functional and structural studies, provided that sufficient CD is added and enough lipids are removed, any tension-sensitive ion channel can be activated by this approach. Given the current interest in eukaryotic MS channels, and that many of these channels respond to membrane forces, CDs are likely an exceptionally useful tool for *in-vitro* studies of this captivating class of membrane proteins.

## Methods

### Lipids

All lipids used in this study, soybean polar azolectin extract (Cat No# 541602), 1,2-dioleoyl-sn-glycero-3-phosphoethanolamine (PE18:1), 1,2-dioleoyl-sn-glycero-3-phosphocholine (PC18:1), 1,2-didecanoyl-sn-glycero-3-phosphocholine (PC10) and 1,2-dioleoyl-sn-glycero-3-phosphoethanolamine-N-(lissamine rhodamine B sulfonyl) (rhodamine-PE18:1), were purchased from Avanti.

### Cyclodextrins

α-cyclodextrin (αCD), β-cyclodextrin (βCD), methyl-β-cyclodextrin (mβCD) and γ-cyclodextrin (γCD) were purchased from Sigma-Aldrich. Saturation of mβCD with cholesterol was carried out as described previously (Christian et al., 1997). Briefly, 100 mg of cholesterol (Sigma-Aldrich) in 1:1 (v:v) chloroform:methanol was added to a glass tube and the solvent was evaporated under a N_2_ stream. 10 ml of 50 mM mβCD was added to the tube and vortexed to release the dried cholesterol from the wall of the tube and then sonicated in a bath sonicator for 5 min. This 100% saturated mβCD:cholesterol solution was incubated on a rotating wheel at 37°C overnight. Immediately before use, the solution was filtered through a 0.45-µm syringe filter (Millipore) to remove excess cholesterol crystals.

### Purification of E. coli MscS and MscL

Wild-type and Gly113Ala mutant *E. coli* MscS were purified as previously reported using an N-terminal 6xHis tag (Zhang et al., 2021). Briefly, MscS was expressed in *E. coli* BL21(DE3) cells, which were grown at 37°C in lysogeny broth medium. When the culture reached an OD_600_ of ∼0.6, protein expression was induced using 1 mM isopropyl β-D-1-thiogalactopyranoside (IPTG). After 4 h at 37°C, cells were harvested by centrifugation at 5,000x g for 10 min. Cells were resuspended in 30 mM Tris-HCl, pH 7.5, 250 mM NaCl, 1% v/v Triton-X100, supplemented with one tablet of cOmplete protease inhibitor cocktail (Sigma-Aldrich) and lysed by sonication. The lysate was centrifuged at 16,000x g for 30 min, incubated with 2 ml nickel resin (Qiagen) and washed with 40 mM imidazole in 30 mM Tris-HCl, pH 7.5, 250 mM NaCl, 0.02% (w/v) dodecyl maltoside (DDM). Protein was eluted with 250 mM imidazole in 30 mM Tris-HCl, pH 7.5, 150 mM NaCl, 0.02% DDM, concentrated using Amicon Ultra 15-ml 50-kDa centrifugal filter unit (Millipore) and loaded onto a Superdex200 column (GE Healthcare) in 30 mM Tris-HCl, pH 7.5, 150 mM NaCl, 0.02% DDM. Fractions containing MscS were pooled and used immediately for reconstitution into nanodiscs or for reconstitution and patch-clamping. For electrophysiological studies, wild-type and Gly22Ser mutant MscL with a 6xHis-tag were purified as previously reported (Rosholm et al., 2017). Briefly, wild-type and Gly22Ser mutant MscL were expressed in *E. coli* BL21(DE3) cells (Novagen), grown at 37°C in lysogeny broth to an OD_600_ of ∼0.8, and then induced with 1□mM IPTG. After 3□h, the cells were centrifuged, the pellet resuspended in phosphate-buffered saline (PBS; 10 mM Na_2_HPO_4_, 1.8 mM KH_2_PO_4_, pH 7.5, 137 mM NaCl, 2.7 mM KCl) with ∼0.02□mg/ml DNase (Sigma DN25) and 0.02% (w/v) PMSF (Amresco M145), and the cells broken with a TS5/48/AE/6□A cell disrupter (Constant Systems) at 31,000□psi at 4°C. Cell debris was removed by centrifugation (12,000x□g for 15□min at 4°C) and membranes were then pelleted at 45,000 rpm in a Type 45 Ti rotor (Beckman) for 3□h at 4°C. Membrane pellets were solubilized overnight at 4°C with 8 mM DDM in PBS, pH 7.5. After centrifugation at 12,000x g for 20□min at 4°C, the supernatant was applied to cobalt sepharose (Talon^®^, 635502, Clontech). The column was washed four times with 40 ml of 35 mM imidazole in PBS, pH 7.5, and protein was eluted using 500 mM imidazole in PBS, pH 7.5. The imidazole concentration was decreased using a 100-kDa Amicon-15 centrifugal filter unit (Merck Millipore) with 1 mM DDM in PBS, pH 7.5.

For structural studies, *E. coli* MscL-pET15b plasmid was purchased from Addgene (Addgene plasmid # 92418; http://n2t.net/addgene:92418; RRID:Addgene_92418). The plasmid with an N-terminal 6xHis tag was used to transform *E. coli* BL21(DE3) cells, which were grown at 37°C in lysogeny broth medium containing 100 µg/ml ampicillin. When the culture reached an OD_600_ of ∼0.6, protein expression was induced by adding IPTG to a final concentration of 1 mM. After another 4 h at 37°C, cells were harvested by centrifugation at 5,000x g for 10 min. Cells were resuspended and lysed by sonication in buffer containing 1% Triton-X100 in 30 mM Tris-HCl, pH 7.5, 250 mM NaCl, supplemented with one tablet of cOmplete protease inhibitor cocktail (Sigma-Aldrich). The lysate was clarified by centrifugation at 16,000x g for 30 min at 4°C, incubated with 2 ml nickel resin (Qiagen) and washed with 40 bead volumes of 40 mM imidazole in 30 mM Tris-HCl, pH 7.5, 250 mM NaCl, 0.02% DDM. Protein was eluted with 250 mM imidazole in 30 mM Tris-HCl, pH 7.5, 150 mM NaCl, 0.02% DDM, concentrated using Amicon Ultra 15-ml 10-kDa centrifugal filters (MilliporeSigma), and loaded onto a Superdex200 column (GE Healthcare) in 30 mM Tris-HCl, pH 7.5, 150 mM NaCl, 0.02% DDM. Fractions containing MscL were pooled and used immediately for reconstitution into nanodiscs.

The membrane-scaffold protein MSP1E3D1, which assembles nanodiscs of 13 nm diameter, with a tobacco etch virus (TEV) protease-cleavable N-terminal 6xHis tag was expressed in *E. coli* BL21(DE3) cells as described for MscL. The cells were lysed by sonication with 1% Triton-X100 in 30 mM Tris-HCl, pH 7.5, 500 mM NaCl, supplemented with one tablet of cOmplete protease inhibitor cocktail (Sigma-Aldrich). After centrifugation at 16,000x g for 30 min at 4°C, the supernatant was loaded on a nickel column, and the beads were washed with 20 column volumes (CV) of 40 mM imidazole in 30 mM Tris-HCl, pH 7.5, 500 mM NaCl, 1% (w/v) sodium cholate, followed by 20 CV of the same buffer without sodium cholate. Protein was eluted with 250 mM imidazole in 30 mM Tris-HCl, pH 7.5, 150 mM NaCl. The His tag was removed by incubation with TEV protease at a molar MSP1E3D1:TEV protease ratio of 30:1. After dialysis overnight at 4°C against 400 ml of 30 mM Tris-HCl, pH 7.5, 150 mM NaCl, the sample was loaded onto a nickel column to remove the cleaved-off His tag and the His-tagged TEV protease. The flow-through was concentrated using Amicon Ultra 15-ml 10-kDa centrifugal filters (MilliporeSigma) and loaded onto a Supderdex200 column equilibrated with 30 mM Tris-HCl, pH 7.5, 150 mM NaCl. The MSP1E3D1-containing fractions were pooled and concentrated to 4.2 mg/ml using Amicon Ultra 15-ml 10-kDa centrifugal filters (MilliporeSigma).

### Reconstitution of MscL into nanodiscs

PC18:1 was solubilized with 20 mM sodium cholate in 30 mM Tris-HCl, pH 7.5, 150 mM NaCl with sonication. MscL and MSP1E3D1 were mixed with the detergent-solubilized lipid at a molar ratio of 1:10:1000 in 12 ml of 30 mM Tris-HCl, pH 7.5, 150 mM NaCl, 0.02% DDM. After 10 min, 1.5 ml of BioBeads SM-2 slurry (Bio-Rad) was added to remove the detergents. After overnight incubation with constant rotation, the BioBeads were allowed to settle by gravity. The supernatant was loaded onto a nickel column to remove the empty nanodiscs. The column was washed with 20 CV of 40 mM imidazole in 30 mM Tris-HCl, pH 7.5, 150 mM NaCl and MscL reconstituted into nanodiscs was eluted with 250 mM imidazole in 30 mM Tris-HCl, pH 7.5, 150 mM NaCl. Samples were concentrated using Amicon Ultra 15-ml 10-kDa centrifugal filters (MilliporeSigma) and loaded onto a Superdex200 column equilibrated with 30 mM Tris-HCl, pH 7.5, 150 mM NaCl. Peak fractions containing MscL in nanodiscs were pooled and used to prepare vitrified samples for cryo-EM.

### Treatment of MscL-containing nanodiscs with β-cyclodextrin (βCD)

PC18:1 nanodiscs containing MscL were incubated with 100 mM βCD as described (Zhang et al., 2021). After 16 h, the sample was loaded onto a Superdex200 column equilibrated with 30 mM Tris-HCl, pH 7.5, 150 mM NaCl. The peak fractions were pooled, concentrated using Amicon Ultra 4-ml 10-kDa centrifugal filters (MilliporeSigma) and used immediately for cryo-EM sample preparation.

### EM specimen preparation and data collection

The homogeneity of samples was first examined by negative-stain EM with 0.7% (w/v) uranyl formate as described (Ohi et al., 2004). The protein concentration was measured with a nanodrop spectrophotometer (Thermo Fisher Scientific) and adjusted to 0.2 mg/ml. Aliquots of 4 µl were applied to glow-discharged 300 mesh R1.2/1.3 Au grids (Quantifoil) using a Vitrobot Mark VI (Thermo Fisher Scientific) set at 4°C and 100% humidity. After 5 s, grids were blotted for 5 s with a blot force of -2 and plunged into liquid nitrogen-cooled ethane.

Cryo-EM imaging was performed in the Cryo-EM Resource Center at the Rockefeller University using SerialEM (Mastronarde, 2005). The data of MscL in PC18:1 nanodiscs before and after treatment with βCD were collected on a 300-kV Titan Krios electron microscope at a nominal magnification of 28,000x, corresponding to a calibrated pixel size of 1.0 Å on the specimen level. Images were collected at a defocus range of -1.2 to -2.5 μm with a K2 Summit direct electron detector in super-resolution counting mode. The ‘superfast mode’ in SerialEM was used, in which 3×3 holes are exposed using beam tilt and image shift before moving the stage to the next position (Cheng et al., 2018). Exposures of 10 s were dose-fractionated into 40 frames (250 ms per frame) with a dose rate of 6 electrons/pixel/s (∼1.38 electrons/Å^2^/frame), resulting in a total dose of 55 electrons/Å^2^.

### Image Processing

The collected movie stacks were gain-normalized, motion-corrected, dose-weighted, and binned over 2×2 pixels in Motioncorr2 (Zheng et al., 2017). The contrast transfer function (CTF) parameters were determined with CTFFIND4 (Rohou and Grigorieff, 2015) implemented in RELION-3 (Zivanov et al., 2018). Particles were automatically picked with Gautomatch (http://www.mrc-lmb.cam.ac.uk/kzhang/Gautomatch/). 124,656 particles were extracted from 1,828 micrographs of MscL in PC18:1 nanodiscs before βCD treatment, and 1,763,603 particles were extracted from 4,037 micrographs for MscL in PC18:1 nanodiscs after βCD treatment. The particles were normalized and subjected to 2D classification in RELION-3.

### Proteoliposome reconstitution

Prior to reconstitution into liposomes, the 6xHis tag was cleaved from MscS with thrombin. MscS and MscL were reconstituted into liposomes with different lipid components using a modified dehydration/rehydration (D/R) reconstitution method (Häse et al., 1995). Azolectin (soybean polar azolectin extract, Avanti) was dissolved in chloroform and mixed with lipids of interest. Fluorescent rhodamine-PE18:1 was added at 0.1% (w/w) and the lipid mixture was dried under N_2_ flow. The lipid film was then suspended in D/R buffer (5 mM HEPES, pH adjusted to 7.2 using KOH, 200 mM KCl) and vortexed, followed by water bath sonication for 15 min. Protein was added to the lipid mixture at ratios (w/w) of 1:200 for MscS and 1:1000 for MscL. After a 3-h incubation with agitation at room temperature, 200 mg of BioBeads (SM-2, BioRad) were added and the sample incubated for another 3 h at room temperature. Finally, the mixture was centrifuged at 40,000 rpm in a Beckman Type 50.2 Ti rotor for 45 min and the lipid mixture was vacuum desiccated in the dark overnight. The proteoliposomes were rehydrated in D/R buffer overnight before use.

### Electrophysiology

Proteoliposomes (0.5 µl which equates to ∼30-100 ng of protein-lipid mixture) were incubated in patch buffer containing 5 mM HEPES, pH adjusted to 7.2 using KOH, 200 mM KCl, 40 mM MgCl_2_ for 15 min until unilamellar blisters formed on their surface. The patch pipette solution contained a symmetrical ionic solution in all recordings. The single-channel currents were amplified using an Axopatch 200B amplifier (Molecular Devices). The *E. coli* MscS and MscL currents were filtered at 2 kHz and sampled at 10 kHz with a Digidata 1440A using pClamp 10 software. Negative hydrostatic pressure was applied using a high-speed pressure clamp (ALA Sciences).

### Patch fluorometry

Wild-type MscS was added to liposomes at a protein:lipid ratio (w/w) of 1:200 and recorded by imaging the tip of the patch pipette using a confocal microscope (LSM 700; Carl Zeiss) equipped with a water immersion objective lens (×63, NA1.15) and housed within a Faraday cage. The excised proteoliposome patches that consisted of 99.9% (w/w) lipids of interest and 0.1% rhodamine-PE18:1 were excited with a 555-nm laser. Fluorescence images of the membranes were acquired and analyzed with ZEN software (Zeiss). To improve visualization of liposome patches, the pipette tip was bent approximately 28° with a microforge (MF-900; Narishige, Tokyo, Japan) to make it parallel to the bottom face of the recording chamber (Nomura et al., 2012). The diameter of the patch dome at a given negative hydrostatic pressure was measured using the ZEN software. Tension was calculated using LaPlace’s law as previously described (Nomura et al., 2015, Shaikh et al., 2014, Cox et al., 2016).

## Acknowledgments

B.M. is supported by a National Health and Medical Research Council of Australia Principal Research Fellowship (APP1135974). C.D.C. is supported by a New South Wales Health Early-Mid Career Research Fellowship.

