## Supplementary Information for "Cyclodextrins increase membrane tension and are universal activators of mechanosensitive channels"

***Running title:*** Cyclodextrin activation of MS channels

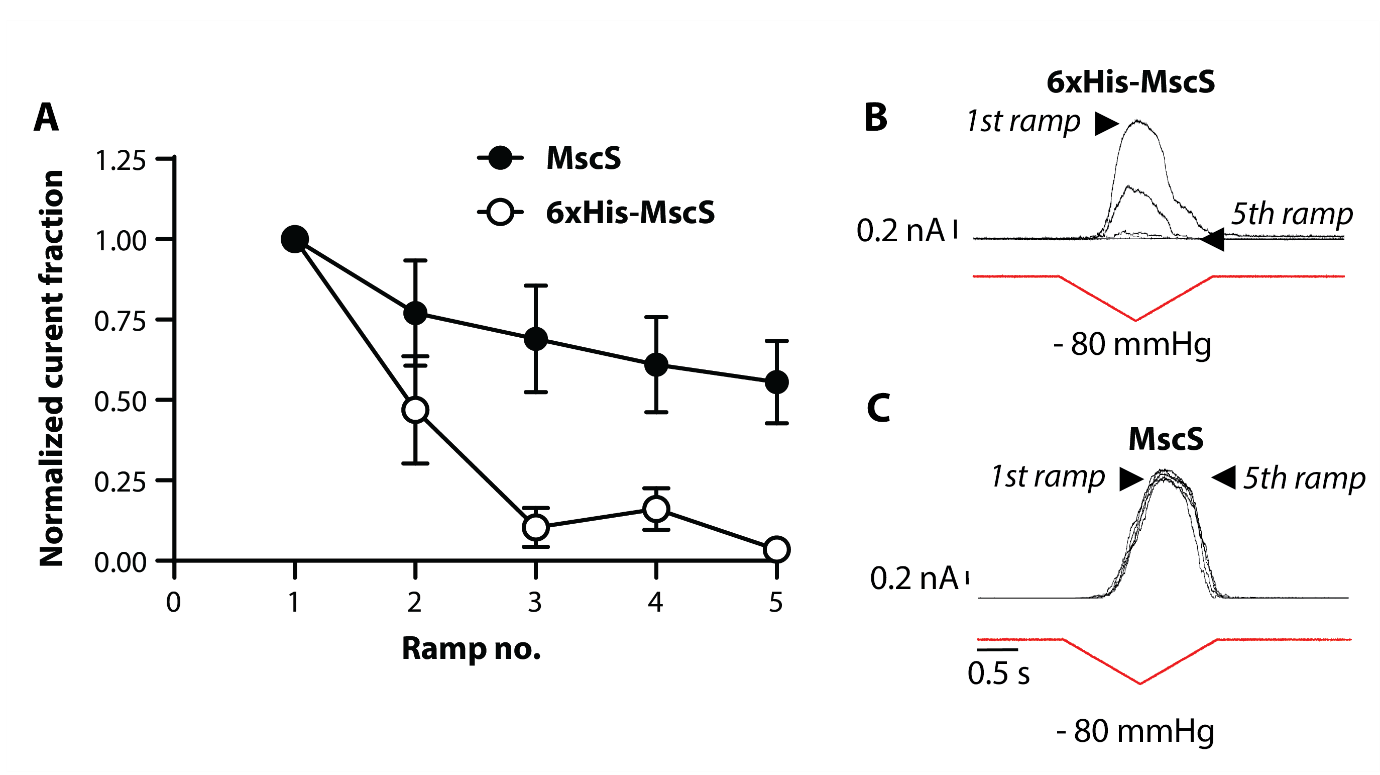
**SI Figure 1. Effect of N-terminal His-tag on azolectin reconstituted MscS currents.** (A) Quantification of normalized current of 6xHis-MscS and MscS with the His-tag cleaved in response to 5 1s long pressure pulses up to a maximum pressure of -80 mmHg (n=5). (B) Representative group of 5 electrophysiological traces showing (B) 6xHis-MscS and (C) MscS with the His-tag cleaved activity in response to pressure in azolectin liposomes.

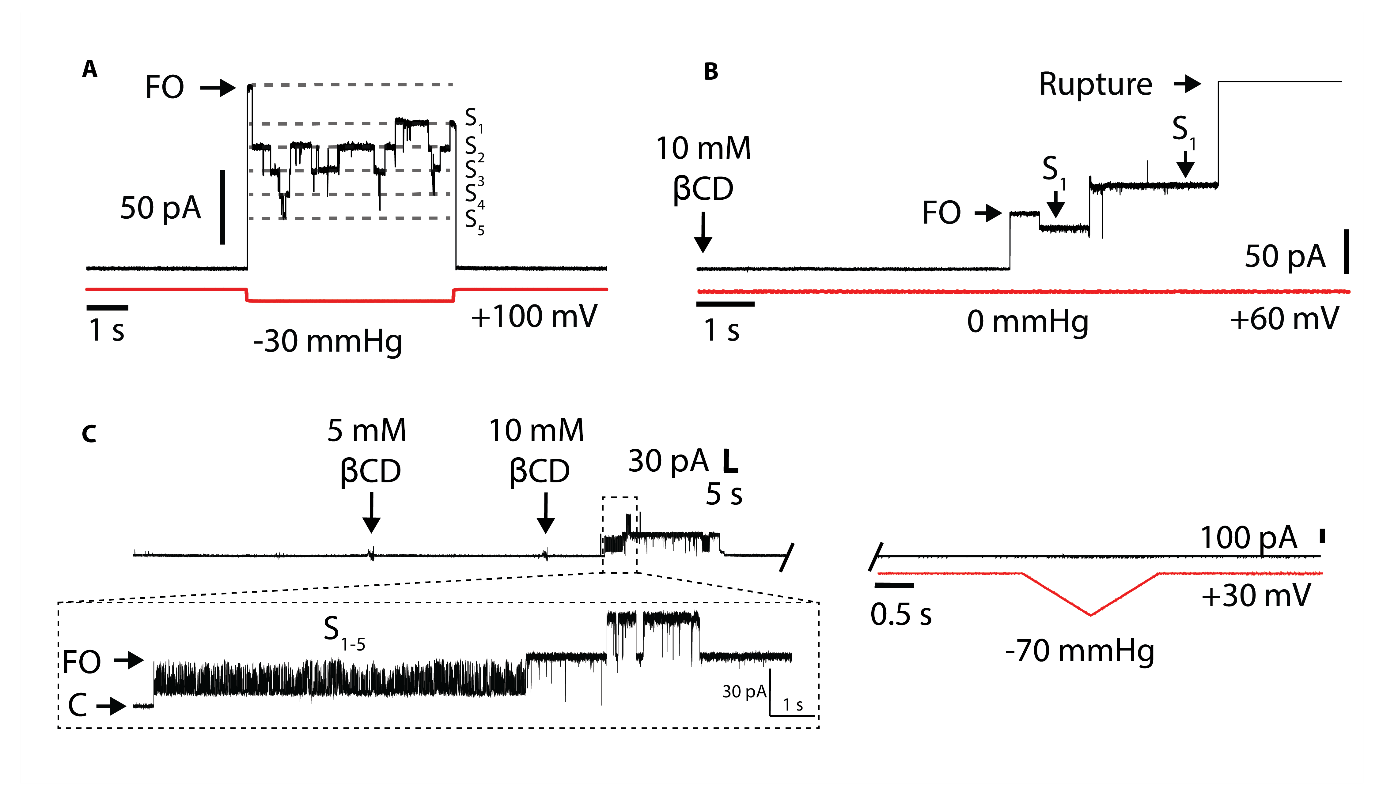

**SI Figure 2. Cyclodextrin treatment causes prolonged sojourns into MscS sub-conducting states**. (A) Representative trace of MscS substate activity in response to pressure at +100 mV pipette potential illustrating multiple long lived substates (S_1-5_) after the fully open channel (FO) is observed. (B) Exemplar trace showing 10 mM βCD can instigate spontaneous opening that also causes substate gating in WT MscS reconstituted into azolectin liposomes. In this trace one prominent substate at ~70 % of the unitary conductance is observed. (C) Exemplar trace showing that 10 mM βCD can instigate gating between many of the WT MscS substates even at modest pipette voltages (+30 mV). This activity then subsides and the channel is desensitized (right panel).

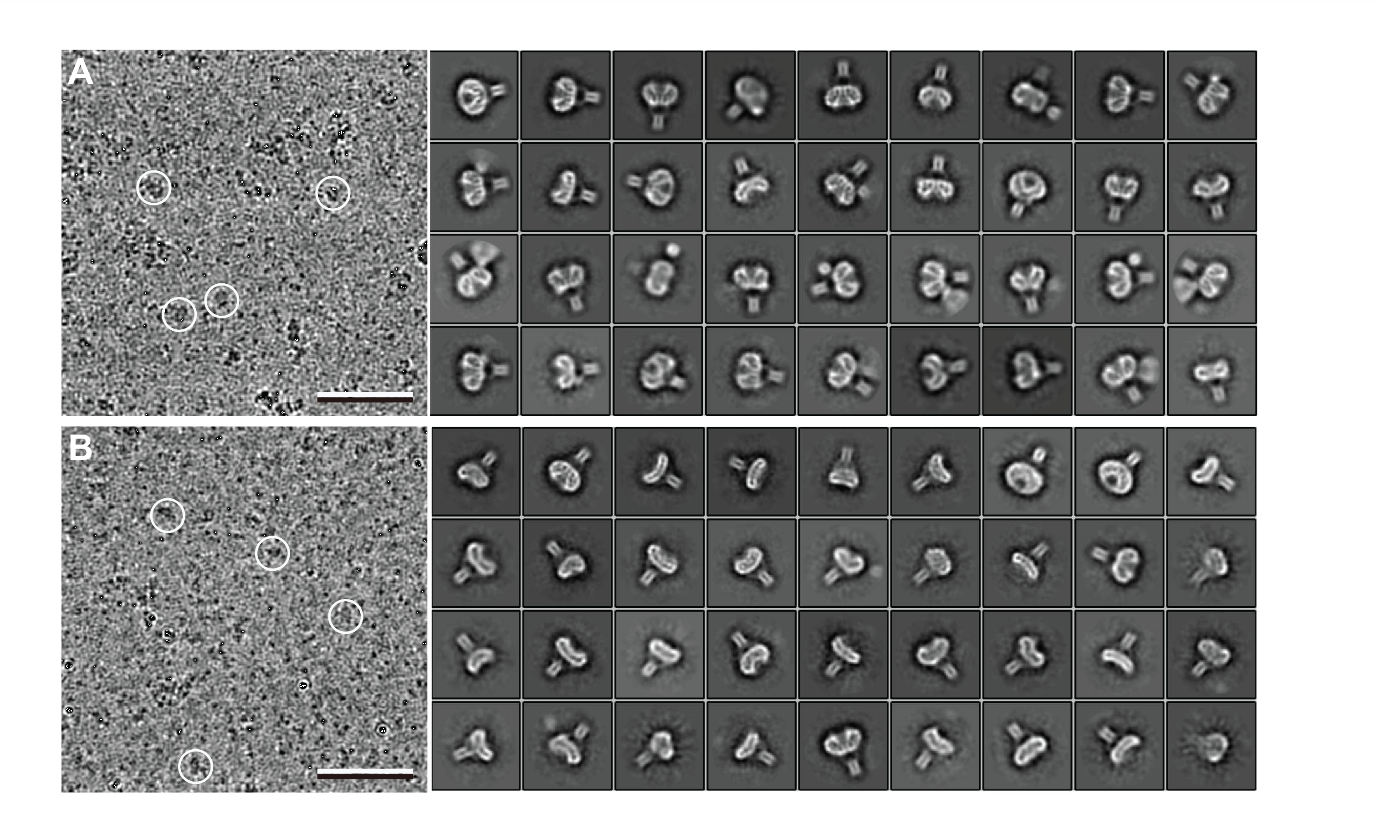

**SI Figure 3. Image processing of MscL in PC18:1 nanodiscs before and after treatment with βCD.** (A) Area of a raw image (left) and selected 2D-class averages (right) of vitrified MscL in PC18:1 nanodiscs before βCD treatment. (B) Area of a raw image (left) and selected 2D-class averages (right) of vitrified MscL in PC18:1 nanodiscs after βCD treatment. Some particles are circled in the raw images. Scale bars, 50 nm. Side length of individual averages, 20 nm.
